# CoBRA: Containerized Bioinformatics workflow for Reproducible ChIP/ATAC-seq Analysis - from differential peak calling to pathway analysis

**DOI:** 10.1101/2020.11.06.367409

**Authors:** Xintao Qiu, Avery S. Feit, Ariel Feiglin, Yingtian Xie, Nikolas Kesten, Len Taing, Joseph Perkins, Ningxuan Zhou, Shengqing Gu, Yihao Li, Paloma Cejas, Rinath Jeselsohn, Myles Brown, X. Shirley Liu, Henry W. Long

## Abstract

ChIP-seq and ATAC-seq have become essential technologies used as effective methods of measuring protein-DNA interactions and chromatin accessibility. However, there is a need for a scalable and reproducible pipeline that incorporates correct normalization between samples, adjustment of copy number variations, and integration of new downstream analysis tools. Here we present CoBRA, a modularized computational workflow which quantifies ChIP and ATAC-seq peak regions and performs unsupervised and supervised analysis. CoBRA provides a comprehensive state-of-the-art ChIP and ATAC-seq analysis pipeline that is usable by scientists with limited computational experience. This enables researchers to gain rapid insight into protein-DNA interactions and chromatin accessibility through sample clustering, differential peak calling, motif enrichment, comparison of sites to a reference DB and pathway analysis.

Code availability: https://bitbucket.org/cfce/cobra

## Introduction

ChIP-seq and ATAC-seq have become essential components of epigenetic analysis. They are employed extensively in the study of protein-DNA interactions and chromatin accessibility respectively. Chromatin immunoprecipitation sequencing (ChIP-seq) is a high-throughput technology that provides unique insights into protein function by mapping genome wide binding sites of DNA-associated proteins. Further, Assay for Transposase-Accessible Chromatin sequencing (ATAC-seq) is a high-throughput technology that is imperative in the assessment of genome-wide chromatin accessibility. While numerous pipelines for analyzing ChIP-seq and ATAC-seq data have been reported in the literature [1–8] there remains a strong need for pipelines that can be run by users who have less experience utilizing computational biology tools. Comparisons between ChIP and ATAC-seq experiments can provide insight into differences in protein occupancy, histone marks and chromatin accessibility (Figure 1a), however, existing analysis pipelines can lack useful components necessary in the analysis. There is a need for better normalization between samples, adjustment of copy number variations, integrating new downstream analysis tools such as cistromeDB Toolkit [9] and integrating epigenetic data with RNA-seq.

**Figure 1.**
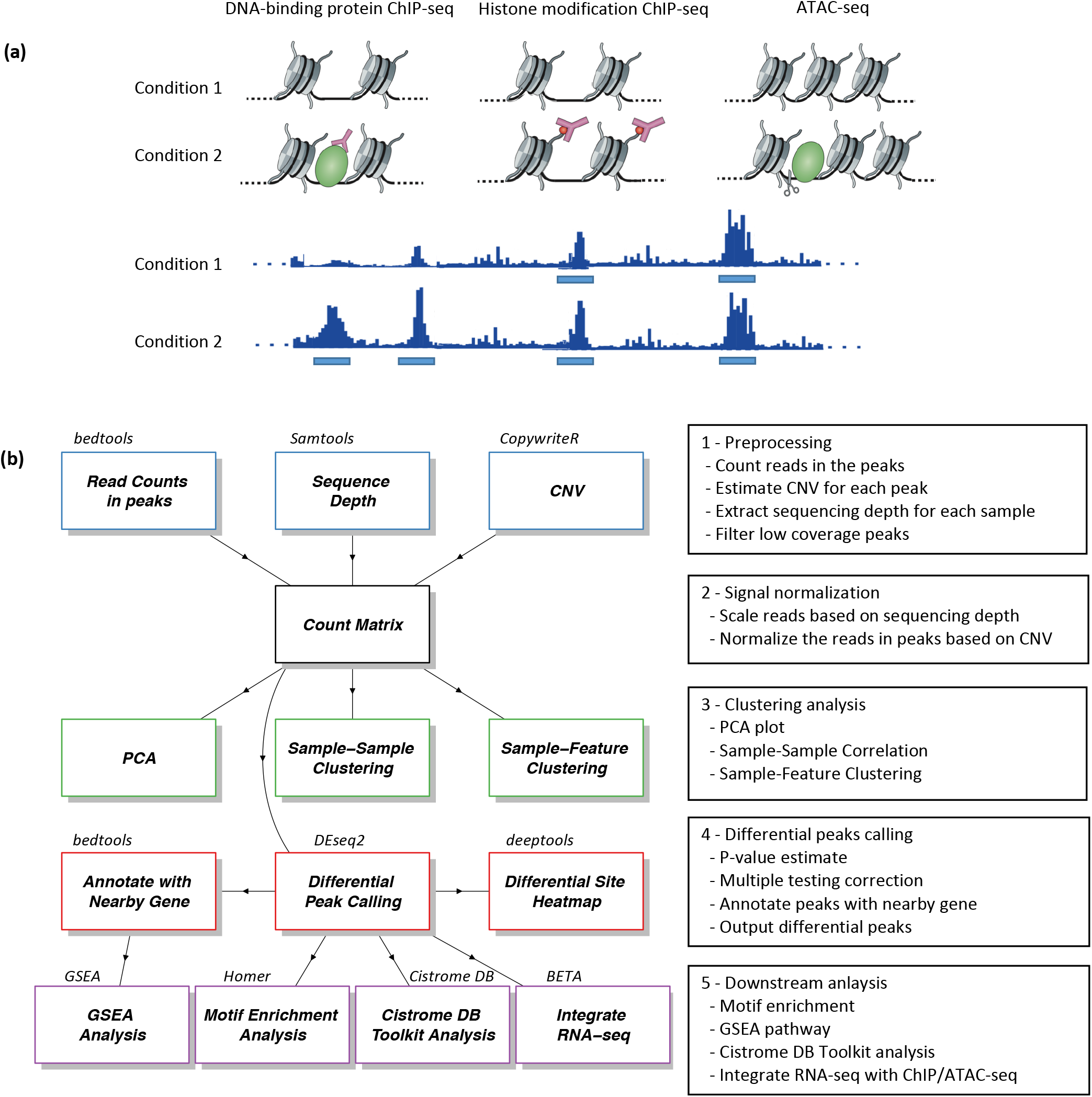
Overview of CoBRA. a) Biological motivation of CoBRA. Comparisons between ChIP and ATAC-seq peaks in well-designed experiments can provide insight into differences in protein occupancy, histone marks and chromatin accessibility. b) Overview of the workflow performed by CoBRA. Read counts are quantified and normalized for sequencing depth and CNV for clustering and differential peak calling analysis. The result of differential peak calling is used downstream for motif enrichment, GSEA, CistroneDB toolkit, and BETA analysis.

In this work we developed CoBRA (Containerized Bioinformatics workflow for Reproducible ChIP/ATAC-seq Analysis), a modularized computational workflow which quantifies ChIP and ATAC-seq peak regions and performs unsupervised and supervised analysis. The pipeline provides sample clustering, differential peak calling, motif enrichment and clustering, comparison of sites to a reference DB and pathway analysis. In addition, it provides clear, high-quality visualizations for all results.

CoBRA uses Snakemake [10], a workflow management system to create the computational pipeline. Using Snakemake system enables the reproducibility and scalability of CoBRA. This framework allows for the addition or replacement of analysis tools as well as for the parallelization of computationally intensive processes. To make CoBRA portable, the workflow and its software dependencies are available as a Docker container, which can be used on any machine with Docker installed. This includes local servers, high-performance clusters, and cloud-based machines. Docker will automatically download all required software dependencies because the container encapsulates all of the supporting software and libraries, eliminating the possibility of conflicting dependencies.

CoBRA therefore provides solutions to challenges inherent in many bioinformatics workflows: it is portable, reproducible, scalable and easy to use. It is open source (https://bitbucket.org/cfce/cobra) and well documented online including step-by-step tutorials that go through all three case studies presented in this paper (https://cfce-cobra.readthedocs.io). This combination of features enables researchers to gain rapid insight into protein-DNA interactions and chromatin accessibility with comprehensive state-of-the-art ChIP and ATAC-seq analysis.

## Methods

### Overall design

The CoBRA pipeline is implemented using the snakemake workflow management system [10] and is described via a human-readable, Python-based language. This allows CoBRA to scale to server, cluster, grid and cloud environments, without the need to modify the workflow. For ChIP-seq and ATAC-seq experiments, CoBRA provides both unsupervised and supervised analyses (Figure 1b). It does not include preprocessing ChIP and ATAC-seq quality control steps as this is best handled within other, specialized pipelines [11].

Further, CoBRA is distributed as a Docker container, which can be used on any machine as long as Docker is installed. Docker containers provide a tool for packaging bioinformatics software. It encapsulates all of the supporting software and libraries, eliminates the possibility of conflicting dependencies, and facilitates the installation of required software. With the built-in snakemake reference rule, CoBRA automatically downloads all needed reference files if they have not been downloaded. As a result, CoBRA is reproducible, portable and easy to deploy. Users specify analysis parameters in a simple human readable configuration file (Supplementary Fig1b). A separate file contains metadata about the samples being analyzed (cell line, treatment, time point, etc.) as well as a specification of the differential comparisons to be performed by the pipeline. This metadata file is in CSV format and can be easily modified in any standard text editor or Excel.

### Unsupervised analysis

The pipeline calculates the Reads per Kilobase per Million Mapped Reads (RPKM) using bed files and bam files provided by the user to normalize for sequencing depth and peak size. The RPKM table is filtered through the removal of sites that have low RPKM across multiple samples. Quantile Normalization (default), Z-score, Log transform are available options to normalize the count matrix. To visualize the similarities between samples in the experiment, sample-sample correlation, principal component analysis (PCA) and sample-feature plot are automatically generated by the pipeline.

The sample-sample correlation plot illustrates the similarity between all of the samples on a pairwise basis. It also provides the clustering result based on the Pearson correlation coefficient, r, where distance = 1 - r. The user can opt for using spearman correlation as well as selecting other distance methods (euclidean, manhattan, canberra, binary, maximum, or minkowski) by simply changing the configuration file. The resulting correlation plot helps to determine whether the different sample types can be separated, i.e., samples of different conditions are expected to be more dissimilar to each other than replicates within the same condition. User provided metadata are used to automatically annotate samples in all unsupervised plots.

Further, CoBRA will produce a principal component analysis (PCA) plot depicting how samples are separated in the first two principal components (those with the largest variance) and samples will be automatically color-coded by all user provided annotations. The PCA plot helps the user to determine if any patterns exist between the samples and if outliers are present. Finally, CoBRA will generate a Sample-Feature heatmap. The heatmap illustrates the clustering of samples based on correlation on the horizontal axis and clustering of peaks on the vertical axis. Peaks on the vertical axis can be clustered by hierarchical or k-means clustering. The sample-feature heatmap elucidates patterns of peaks across samples and identifies the clusters that are enriched in a subset of samples.

### Supervised analysis

A common question being asked is what the differential sites are (TF binding/ histone modification/ chromatin accessibility) between sample groups. Several tools (DEseq2, edgeR, Limma) currently available can be applied to analyze differential sites, most of which are derived from RNA-seq count analysis. However, there are differences between the RNA-seq and ChIP-seq count analysis. In RNA-seq experiments, most reads are in the exome, where reads can be normalized by the total number of reads mapped to all genes. In contrast, most ChIP-seq reads are outside of peaks. The FRiP score (fraction of reads in peaks) typically ranges from 1 to 40 percent [11]. Reads in peaks are only a portion of total reads that have been sequenced. Therefore, all reads need to be normalized by the total number of uniquely mapped reads to account for sequence depth. CoBRA uses the bam file to calculate sequencing depth. It utilizes sequencing depth as a scale factor in differential peak calling by DESeq2 (although the user can specify reads in peaks for scaling if specifically required). This is an essential step in differential peak calling. The default scale factor utilized by DESeq2 to normalize the data is the total number of reads mapped to peaks. This method can result in the calling of false positive differential peaks. Using sequencing depth as the scale factor ensures that reads are normalized for experimental variation and not biological variation between samples.

Multiple comparisons can be done within a single run. For each comparison, the number of differential peaks for two adjusted p-value cutoffs and two fold-change cut-offs will be displayed in a summary chart. Further, the bigwig files are used to plot the peak intensity of the differential peaks in a heatmap using deepTools2 [12].

The differentially enriched regions from DEseq2 for each comparison are subsequently run through HOMER [13] for motif enrichment analysis. Motif enrichment analysis is a fundamental approach to look for transcription factor motifs that might be enriched in peaks of interest. We use Homer in the pipeline to look for known and de novo motifs that are enriched in the differential peak regions compared to GC matched, randomly selected genome background. In addition, we utilize a motif clustering algorithm to organize various motifs by similarity making the output (Supplementary Figure 3) easier to evaluate for distinct results. By mapping the peaks to the nearest gene, CoBRA uses GSEA pre-ranked analysis to investigate the pathways that are enriched and depleted for both up and down peaks.

The up and down-regulated sites are also automatically compared to a comprehensive database of ChIP/ATAC and DNase data [9; 14]. This Cistrome Toolkit analysis determines the most similar samples in terms of genomic interval overlaps with the differential sites. The toolkit is particularly useful to identify the major transcription factors related to the differential perturbations. In addition, it can be useful in the identification of the potential biological source (cell line, cell type and tissue type) of the regions of interest.

## Results

In order to illustrate the utility of CoBRA, we applied it to three projects with components that illustrate the different capabilities of our workflow: a GR ChIP-seq data set from the ENCODE project, H3K27ac ChIP-seq data from colon cancer cell lines, and an ATAC-seq experiment on HL-60 promyelocytes differentiating into macrophages. Each example demonstrates some key functions of the CoBRA pipeline.

### Case Studies

#### Example 1: Normalizing GR ChIP-seq data in a dose-response experiment

We downloaded publicly available glucocorticoid receptor (GR) ChIP-seq data (<u>GSE32465</u>) from a lung adenocarcinoma cell line (A549) at 3 different concentrations of dexamethasone, a potent GR agonist. In an analysis of this dataset [15], it was found that the number of Glucocorticoid Receptor (GR) binding sites increases with increasing dexamethasone concentration. In the experiment, samples were treated with 0.5nM, 5nM, or 50nM of dexamethasone. CoBRA’s unsupervised analysis showed that the sample replicates cluster tightly together. Similarities and differences between samples are illustrated by the correlation between treatments vs within treatment in the dendrogram at the top of sample-sample heatmap (Figure 2a), as well as the principal component plot (Supplementary Figure 1).

**Figure 2.**
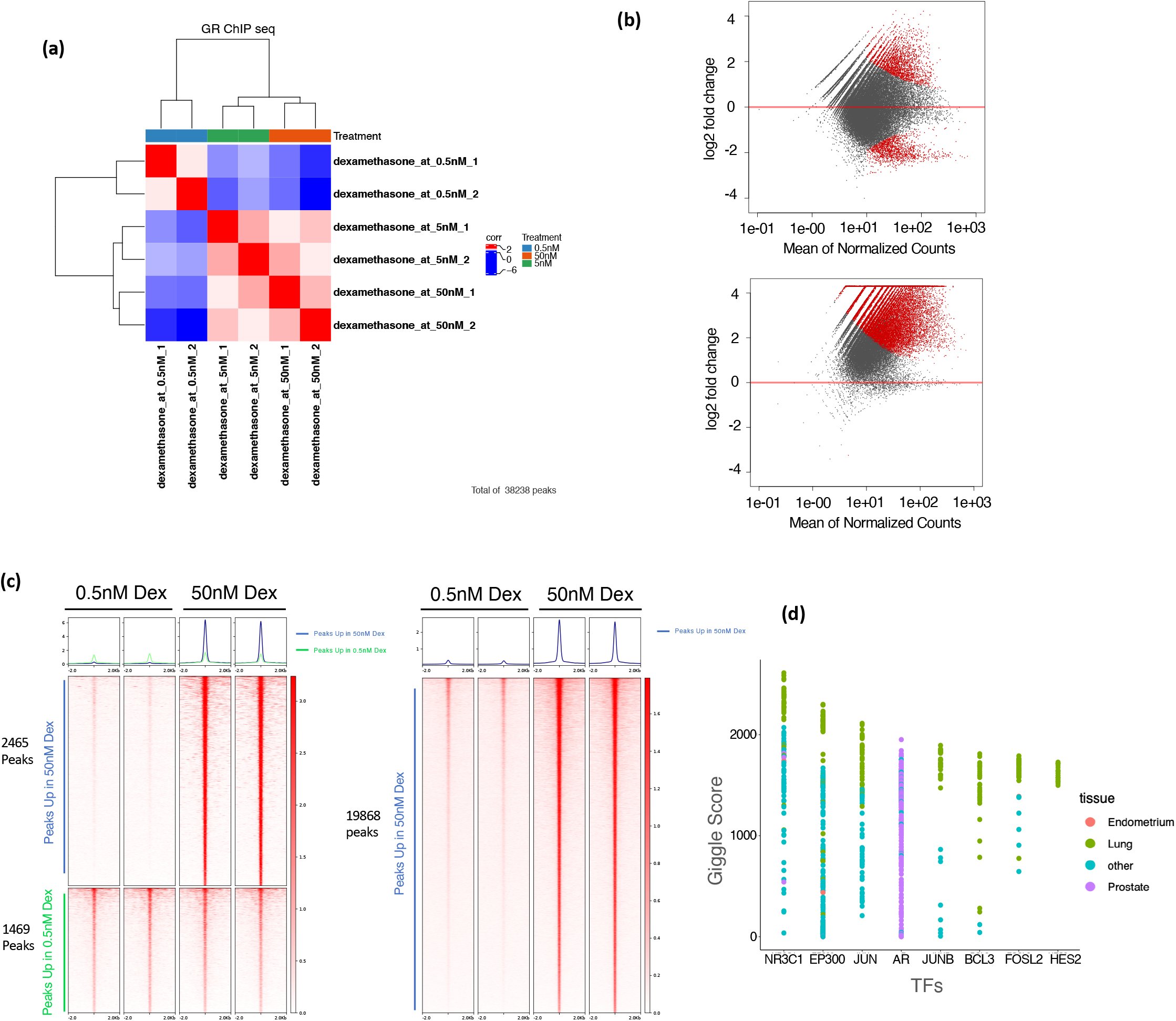
Example of unsupervised and supervised analysis of differential GR binding in A549 cells. a) Sample-Sample heatmap depicting clustering and correlation between A549 cells treated with varying concentrations of dexamethasone. b) Visualization the differences in the GR binding between the 0.5 and 50nM samples, plotted using mean of the peak intensities versus log2(fold change). This illustrates the change in the inferred differential GR binding profile following normalization using scaling factor determined by total reads in peaks (top) and sequencing depth (bottom). c) Deeptools heatmap illustrating differential peaks called by DESeq2 using default scaling factor by total reads in peaks (left) or using scaling factor determined by sequencing depth (right). A group of peaks at the bottom of the left figure exhibit similar binding intensity, however, they are considered downregulated in 50nM treatment in the default DEseq2 result. d) Cistrome Toolkit result illustrating publicly available ChIP seq datasets ranked by binding profile similarity to gained GR binding sites with dexamethasone treatment.

While unsupervised analyses are useful, the advantage of the CoBRA pipeline is its ability to accurately call differential peaks accounting for a variety of factors. We applied DESeq2 to assess the differences in peak binding for samples treated with 50nM of dexamethasone versus samples treated with 0.5nM of dexamethasone. Utilizing DESeq2’s default scale factor method which normalizes the data using the total number of reads in peaks, differential peaks are called (Figure 2b) where they are clearly not present (Figure 2c left). A group of peaks at the bottom of the figure 2c exhibit similar binding intensity, however, they are considered downregulated in 50nM treatment in the DEseq2 result.

DEseq2 by default normalizes all samples by total reads in the read count table. In RNA-seq, most reads are in the exome, where reads can be normalized by the total number of reads mapped to all genes. In contrast, in the GR ChIP-seq experiment, samples treated with 50nM dexamethasone exhibit much greater GR binding and the FRiP score is higher than samples treated with 0.5nM (9.3 vs 0.9). Therefore, DESeq2’s normalization method decreases the peak intensity in the 0.5nM treated samples because the FRiP scores are higher in the 50nM sample resulting in false positive differential peaks (Figure 2c right). In CoBRA, we use a scaling factor dependent on the sequencing depth of each sample. This eliminates the false positive downregulated peaks called by DESeq2 using the default scaling factor (Figure 2c right and Figure 2b). Furthermore, more real differential gained peaks have been successfully identified with CoBRA’s scaling method.

An additional feature of CoBRA is that it automatically analyzes the differential peaks to provide additional insight into their origin and identify similar systems in the literature. In one analysis it determines the most similar ChIP-seq data that is available in a large, curated database of ChIP and ATAC data - cistrome.org [14]. For the gained GR binding sites in the dexamethasone treatment, the result from the Cistrome Toolkit [9] clearly shows that the *NR3C1* in lung tissue is the most similar ChIP-seq in the cistrome database (Figure 2d). CoBRA provides a list of GEO accession numbers corresponding to all ChIP-seq data with similarity to the differential peak set. Using these identifiers, ChIP seq data of interest can be downloaded for further investigation from Cistrome DB[14]. While obviously correct in this simple case, this tool can provide unique insight into gained or lost sites such as identifying which transcription factor potentially binds to a differential peak set after a perturbation and in investigating similar cellular systems. In addition, CoBRA performs a de novo motif analysis on differential sites which can help to identify potential transcriptional regulators enriched in our differentially accessible chromatin elements. In this example the top cluster has all hormone receptor motifs enriched in the upregulated peaks.

#### Example 2: Correcting for Copy Number variation in H3K27ac ChIP-seq

We further illustrate the advantages of the CoBRA pipeline utilizing data from colorectal cancer cell lines. Microsatellite Instable (MSI) and Microsatellite Stable (MSS) are two classes used to characterize colorectal cancers. To analyze these cell lines, we selected six publicly available datasets from several experiments: three MSI samples and three MSS samples [16-20] (<u>GSM1866974</u>, <u>GSM2265670</u>, <u>GSM1224664</u>, <u>GSM1890746</u>, <u>GSM2058027</u>, <u>GSM1890746</u>).

MSS tumors are one of the most highly mutated tumor types [21] and typically exhibit a high number of copy number alterations. Without adjustment, a differential peak caller will rank peak loci with high copy number gain in MSS as being the most differential compared to MSI. These genetic differences, while important, can obscure important epigenetic differences between MSI and MSS. In order to observe differential peaks other than those called as a result of the presence of CNV, copy number variation adjustment was conducted on all samples. For this example the copy number was called using the ChIP-seq data itself with CopywriteR [22] but can also be done with qDNAseq [23] using the input control if available. Any other source of CNV data can also be used when put in a standard igv format. This CNV adjustment alters the differential peaks called by DESeq2. In the case of the MSS vs. MSI comparison, many peaks at the 8q region of the chromosome are being called significantly differential (Figure 3a) but, following CNV correction, the number of differential peaks in this region significantly decreased (Figure 3b).

**Figure 3.**
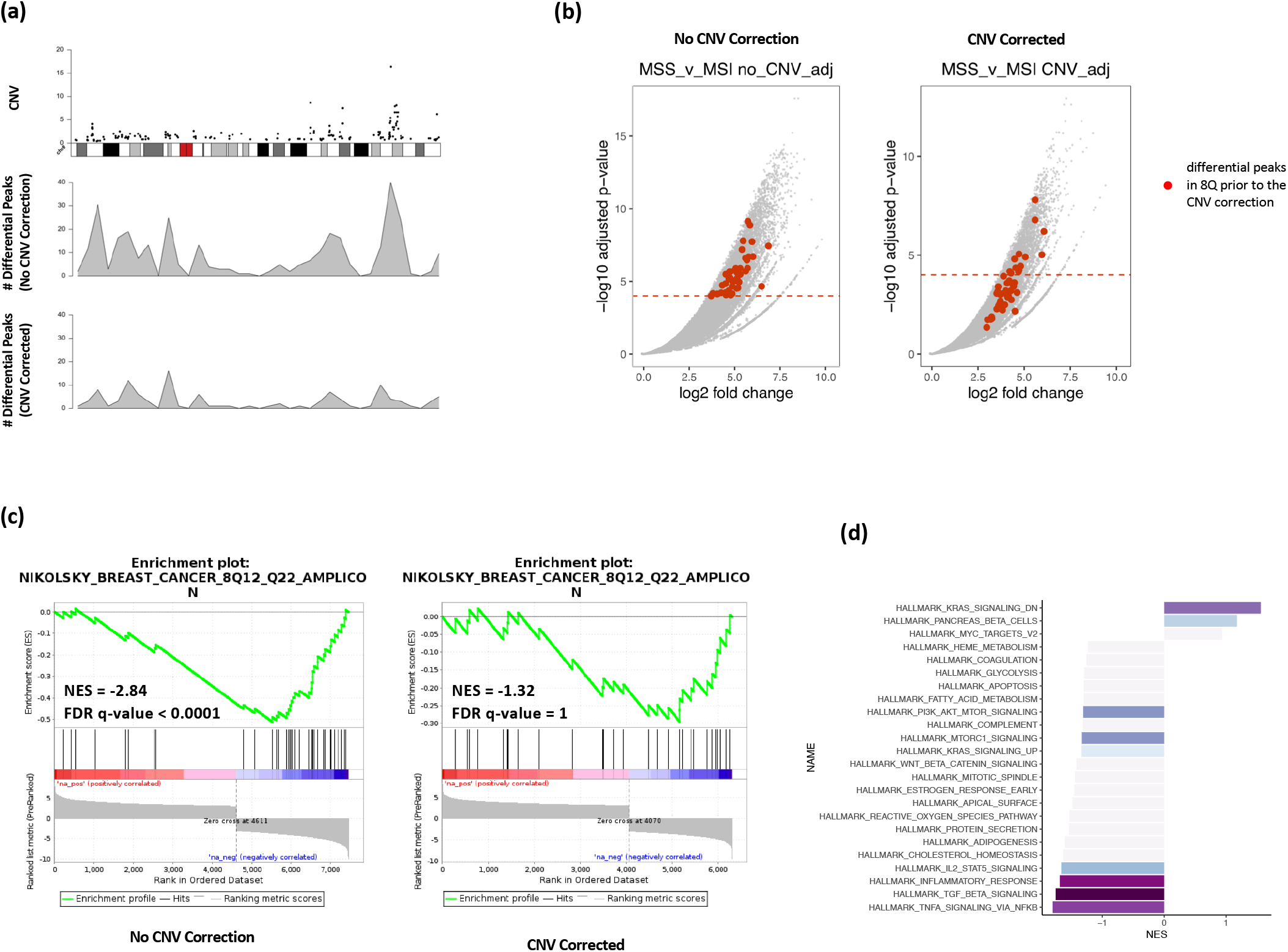
Identification of differential sites correcting for copy number. a) Copy number distribution for an MSS sample on Chromosome 8 (top). Distribution of differentially called peaks with (middle) and without (bottom) CNV adjustment between MSS and MSI cell lines. b) Significant differential peaks in 8Q prior to CNV correction are highlighted. X axis is the log2 fold change and y axis is −log10 of the adjusted p-value. Left side is without CNV correction, and the right side is with CNV correction. c) Enrichment plot for NIKOLSKY_BREAST_CANCER_8Q12_Q22_AMPLICON gene set without (left side) and with (right side) CNV adjustment. d) Enrichment of Hallmarks gene sets after CNV correction based on the differential peak ranking comparing MSS with MSI.

Gene Set Enrichment analysis is performed on the ranked list of genes produced by CoBRA. Without CNV adjustment, GSEA can indicate greatest enrichment in gene sets solely related to amplification. As a result, it is challenging to assess the true epigenetic differences between the two colorectal cancer types. For instance, the gene set ‘NIKOLSKY_BREAST_CANCER_8Q12_Q22_AMPLICON’ includes genes up-regulated in non-metastatic breast cancer tumors with amplification in the 8q22 region. Without adjustment for copy number variation, this gene set is significantly enriched (Figure 3c). It is the 3th ranked gene, with a normalized enrichment score of −2.84 and an adjusted p-value less than 0.0001. With CNV adjustment, this gene set is far less enriched (Fig. 3c). It is the 468th ranked gene set and has a normalized enrichment score of −1.32 and an adjusted p-value of 1.

After CNV correction, the Hallmarks GSEA analysis shows that the MSI cell line has enrichment in the following pathways: TNFA signaling via NFKB, TFG beta signaling, and Inflammatory response (Figure 3d). This is consistent with the literature[24-25] in reference to colon cancer with MSS tumors exhibiting more inflammatory signaling.

#### Example 3: Unsupervised analysis of time series ATAC-seq data

In this example, we illustrate the efficacy of CoBRA’s analysis of ATAC-seq experiments by following the chromatin accessibility profile of differentiating cells [26]. In this experiment researchers utilized a five-day time course (0hr, 3hr, 24hr, 96hr, and 120hr) to profile accessible chromatin of HL-60 promyelocytes differentiating into macrophages (<u>GSE79019</u>). The CoBRA output includes a principal component analysis (PCA) plot (Figure 4a) that demonstrates the temporal differentiation of the macrophages; the early time point is on the left side while the late time point is on the right. Furthermore, the output includes a sample-feature heatmap utilizing k-means (k=3) clustering (Figure 4b) that further illustrates the dramatic differences in open chromatin profiles. The three clusters show clear differences in open chromatin between the early (cluster 1), intermediate (cluster 2), and late stage (cluster 3) time points.

**Figure 4.**
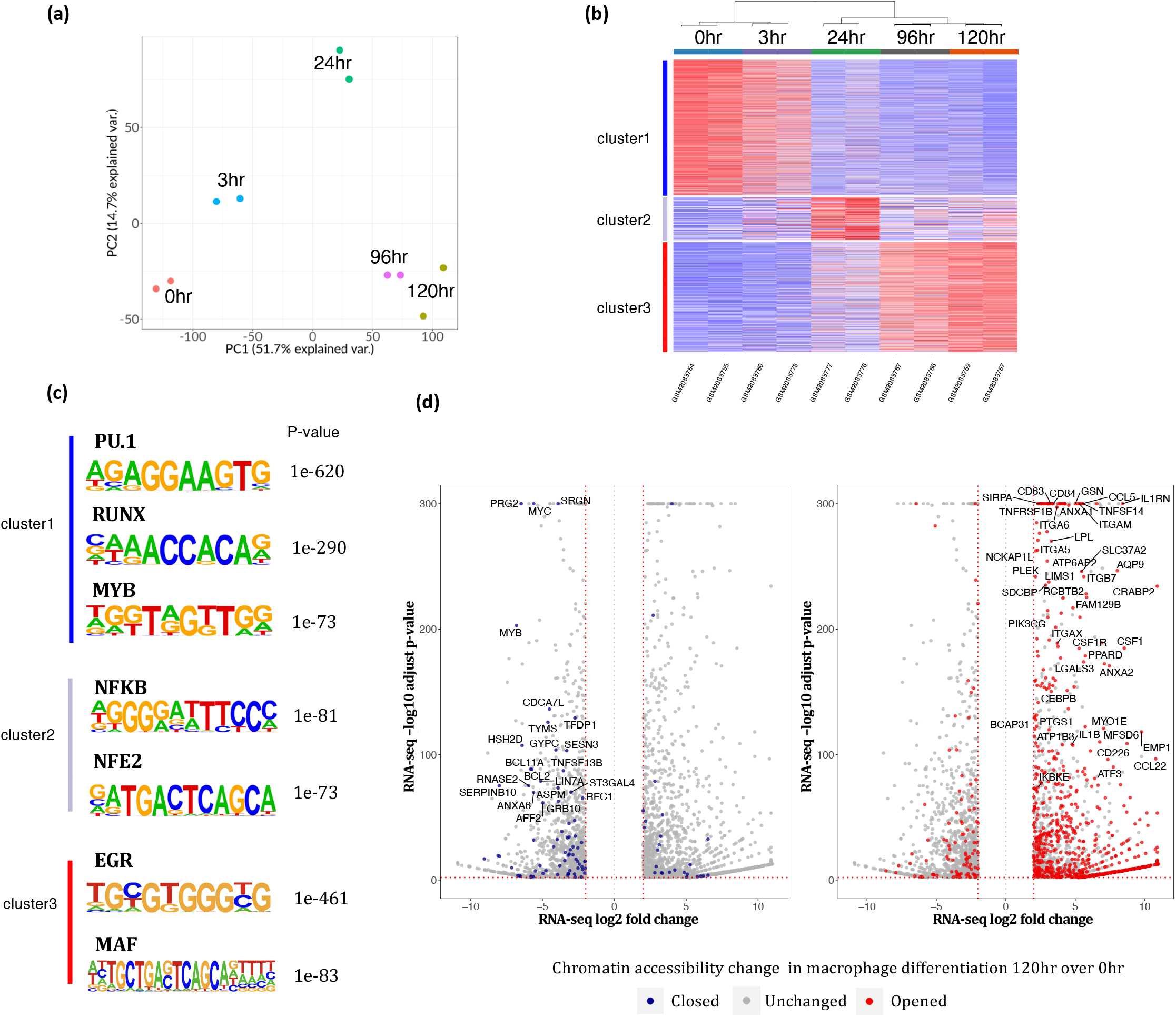
Analysis of ATAC-seq from HL-60 promyelocytes differentiating into macrophages with CoBRA. a) PCA plot depicting how samples cluster along the first two principal axes. b) Sample-Feature heatmap created by CoBRA which depicts sample clustering on the horizontal axis and chromatin accessibility clustering on the vertical axis. Cluster 1,2, and 3 represent sites open at early, middle and late differentiation stages respectively. c) Motifs enriched in early, middle, and late stage differentiation identified by CoBRA. d) Genes differentially expressed during macrophage differentiation (120hr over 0hr). Those genes that also have nearby differential chromatin changes during differentiation are highlighted.

CoBRA automatically performs a de novo motif analysis on each of the three clusters of accessible sites to identify motifs of potential transcriptional regulators enriched in differentially accessible chromatin elements. This analysis identified many transcription factor motifs enriched in each cluster (Figure 4c). Motifs for PU.1, RUNX and MYB were enriched in cluster 1, which exhibits a decrease in accessibility during myeloid differentiation. It is likely that a depletion of PU.1, RUNX and MYB occupancy occurs at these elements during cellular commitment. In addition, we observe the EGR and MAF motifs in clusters 3 suggesting a gain of EGR and MAF occurs at these elements during macrophage differentiation. The motif analysis for cluster 2 also identified chromatin element NFKB and NFE2 as being active during differentiation and depleted in the latter stages. All of these findings are consistent with the results from published papers [26].

Finally, ChIP-seq and ATAC-seq data is often generated in parallel with RNA-seq on the same samples. An extension to CoBRA can take the differential expression gene list from RNA-seq analysis tools such as VIPER [27] and highlight differentially expressed genes that also exhibit differential chromatin accessibility. The volcano plot in Figure 4d is a visualization of the genes differentially expressed during macrophage differentiation and highlights those genes that also have nearby opening chromatin during differentiation. Genes near open chromatin during differentiation are more likely to be upregulated. This profile that combines chromatin accessibility with gene expression can provide insight to potentially identify major transcriptomic elements driving differentiation.

## Discussion

The case studies that we have presented highlight typical use cases for CoBRA. The first example is accurate identification of differential peaks using appropriate normalization of ChIP-seq data. Some methods fail to normalize correctly in calling differential peaks when the FRiP score is impacted by perturbations. CoBRA reduces false positives and identifies more true differential peaks by correctly normalizing for sequencing depth.

The second example demonstrates how CoBRA can be used to account for amplification due to copy number variation present in experimental samples. This is an important feature, as copy number variation can drive the greatest differences between some tumor samples and obscure other biological changes to the cistrome that occur as a result of treatment or other experimental conditions. After adjustment of CNV, differential peaks called by DESeq will not be affected by amplification between samples, allowing biologists to better understand whether differences are caused by changes in the genetic or epigenetic landscape.

The third example illustrates how CoBRA can be applied to ATAC-seq experiments. Unsupervised analyses can identify changes in the chromatin accessibility over time with treatment, and clustering provides insight into similarities and differences between samples and the investigation of the transcription factor motif enrichment in each cluster.

The application of CoBRA to these experiments demonstrate the broad capabilities of the workflow in analyzing ChIP-seq or ATAC-seq experiments. While other workflows used to analyze ChIP or ATAC experiments exist, they lack some of the features present in CoBRA (table 1). Additionally, the highly modular Snakemake framework allows for rapid integration of new approaches or replacement of existing tools. Modules can be added simply by adding a new Snakemake “rule” and adding a flag in the config file (Supplementary Figure 2a–c) to turn the analysis on. Further, CoBRA’s “rules” can be composed of tools written in R, Python, or shell script. The framework allows for great flexibility because each module can be evaluated in its own environment using different tools (e.g. Python 2.7 and Python 3 based software).

**Table 1.** A comparison of the features of CoBRA with other available pipelines.

| Pipelines Features | CoBRA | Diffbind | HMCan-diff | ChIPcomp | Deeptools | esATAC | OPENANNO |
| --- | --- | --- | --- | --- | --- | --- | --- |
| Sample-sample Correlations | ✓ |  | ✓ |  | ✓ | ✓ |  |
| Sample-feature Clustering | ✓ | ✓ |  |  | ✓ |  |  |
| PCA analysis | ✓ | ✓ |  |  |  |  |  |
| Normalize based on Sequencing Depth | ✓ | ✓ |  |  |  |  |  |
| CNV Correction for Differential Peak Calling | ✓ |  | ✓ |  |  |  |  |
| Motif Analysis | ✓ |  |  |  |  | ✓ |  |
| Pathway Analysis | ✓ |  |  |  |  | ✓ |  |
| Package Easy Update | ✓ | ✓ |  | ✓ |  | ✓ | ✓ |
| Easy Support New Species | ✓ |  |  |  |  |  |  |
| Docker Containerized | ✓ |  |  |  |  |  |  |
| Annotate Peak Regions with Public ChIP-seq database | ✓ |  |  |  |  |  | ✓ |
| Step-by-Step Tutorial with Multiple Case Studies | ✓ | ✓ |  |  | ✓ | ✓ |  |

The methods for installing, deploying, and using CoBRA along with a detailed tutorial are provided in the documentation available online(https://cfce-cobra.readthedocs.io/). The workflow was designed to work with Docker, which allows the user to automatically download all required software dependencies, eliminating the possibility of conflicting dependencies. This makes CoBRA easy for those with limited computational training to install and run the workflow. Furthermore, the user does not need to prepare any reference files, as CoBRA automatically downloads all needed reference files. As a result, CoBRA is portable, reproducible and easy to deploy.

In summary we have developed a new pipeline, CoBRA (Containerized Bioinformatics workflow for Reproducible ChIP/ATAC-seq Analysis), that is fast, efficient, portable, customizable and reproducible. The workflow builds upon the ongoing effort to make computational research reproducible using defined workflows running inside Docker containers. CoBRA allows users of varying levels of technical skill to quickly process and analyze new data from ChIP-seq and ATAC-seq experiments. It is the authors’ hope that CoBRA can be a starting point for others to build upon and improve CoBRA as a tool and extend its ability to analyze the cistrome.

## Availability of data and software

The dataset(s) supporting the conclusions of this article are all publicly available in the NCBI Sequence Read Archive as referenced in the text.

The software described in this article is publicly available online.

Project name: CoBRA.

Project home page: https://bitbucket.org/cfce/cobra, https://cfce-cobra.readthedocs.io

Archived version: publication.

Operating system(s): UNIX; MacOS.

Programming language: multiple.

Other requirements: Docker, wget, git, miniconda3.

License: GNU GPL.

Any restrictions to use by non-academics: N/A.

## CRediT author statement

Xintao Qiu: Methodology, Software, validation, Writing - Original Draft

Avery S. Feit: Methodology, Software, validation, Writing - Original Draft

Ariel Feiglin: Methodology, Software, validation

Yingtian Xie: Validation, Data Curation

Nikolas Kesten: Validation

Len Taing: Software, Validation

Joseph Perkins: Software

Ningxuan Zhou: Validation, Investigation

Shengqing Gu: Validation, Investigation

Yihao Li: Validation, Investigation

Paloma Cejas: Validation, Investigation

Rinath Jeselsohn: Resources, Validation

Myles Brown: Conceptualization, Supervision, Funding

X. Shirley Liu: Supervision, Project Administration

Henry W. Long: Supervision, Funding, Conceptualization, Writing - Review & Editing

## Competing Interests

The authors have declared that no competing interests exist.

## Acknowledgements

H.W.L. and M.B. acknowledge funding from the National Institutes of Health (USA) grant 2PO1CA163227.

---

**Supplementary Figure 1.**
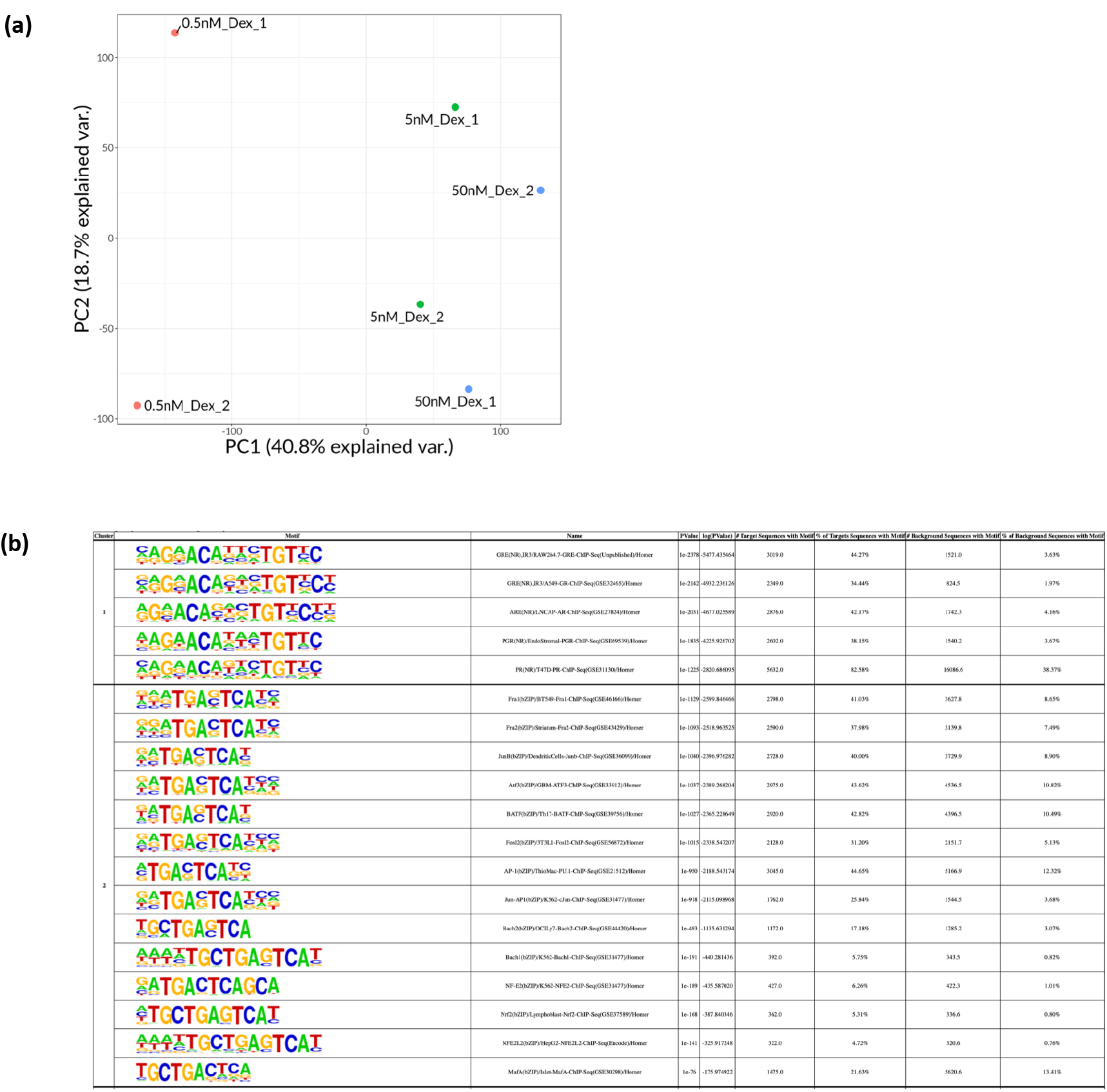
a) PCA plot depicting similarity between dexamethasone treated samples. b) Clustering result of the motif enrichment for sites up with 50nM treatment over 0.5nM.

**Supplementary Figure 2.**
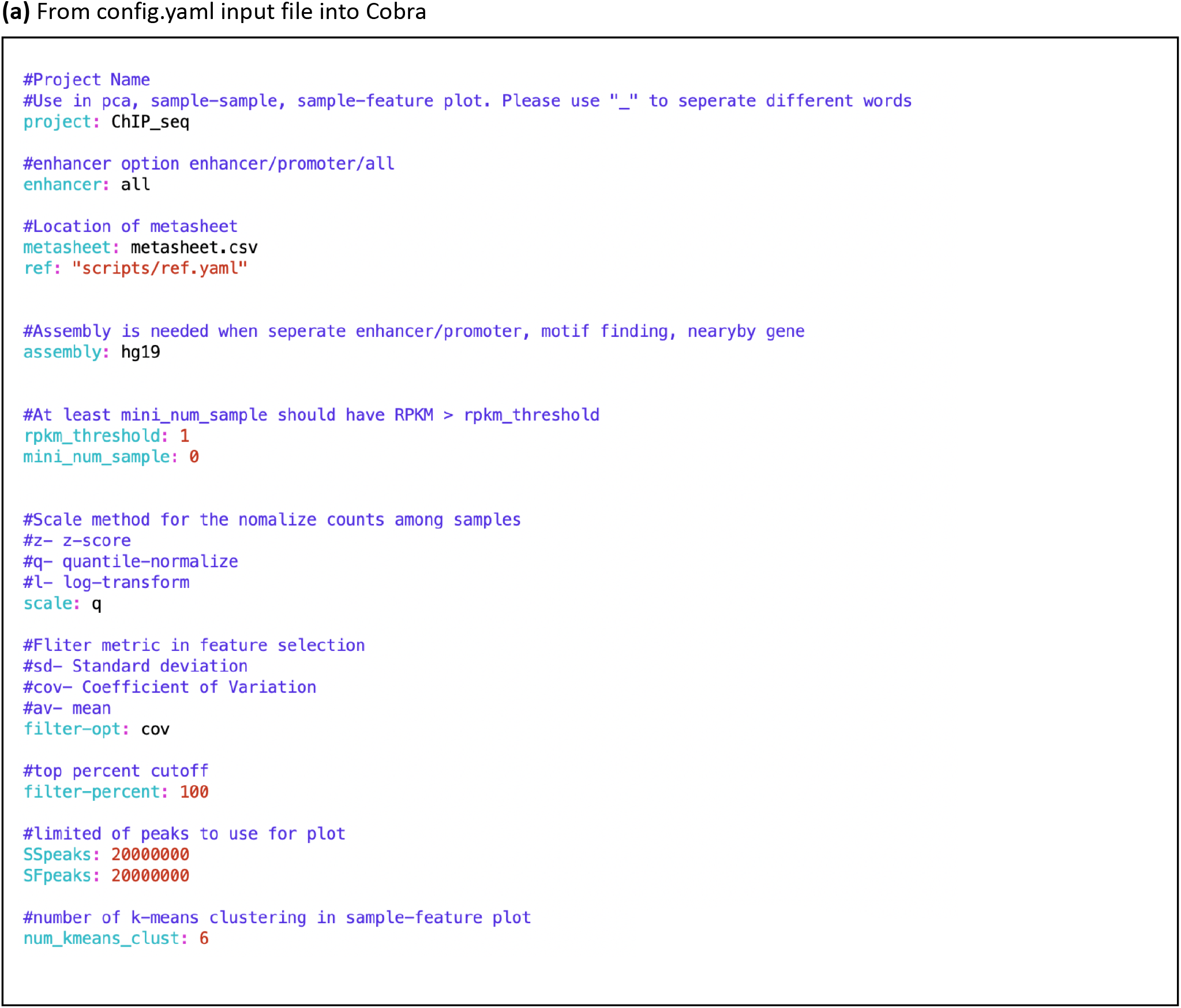
a) Example of parameter setup in config.yaml file.

**Supplementary Figure 2b.**
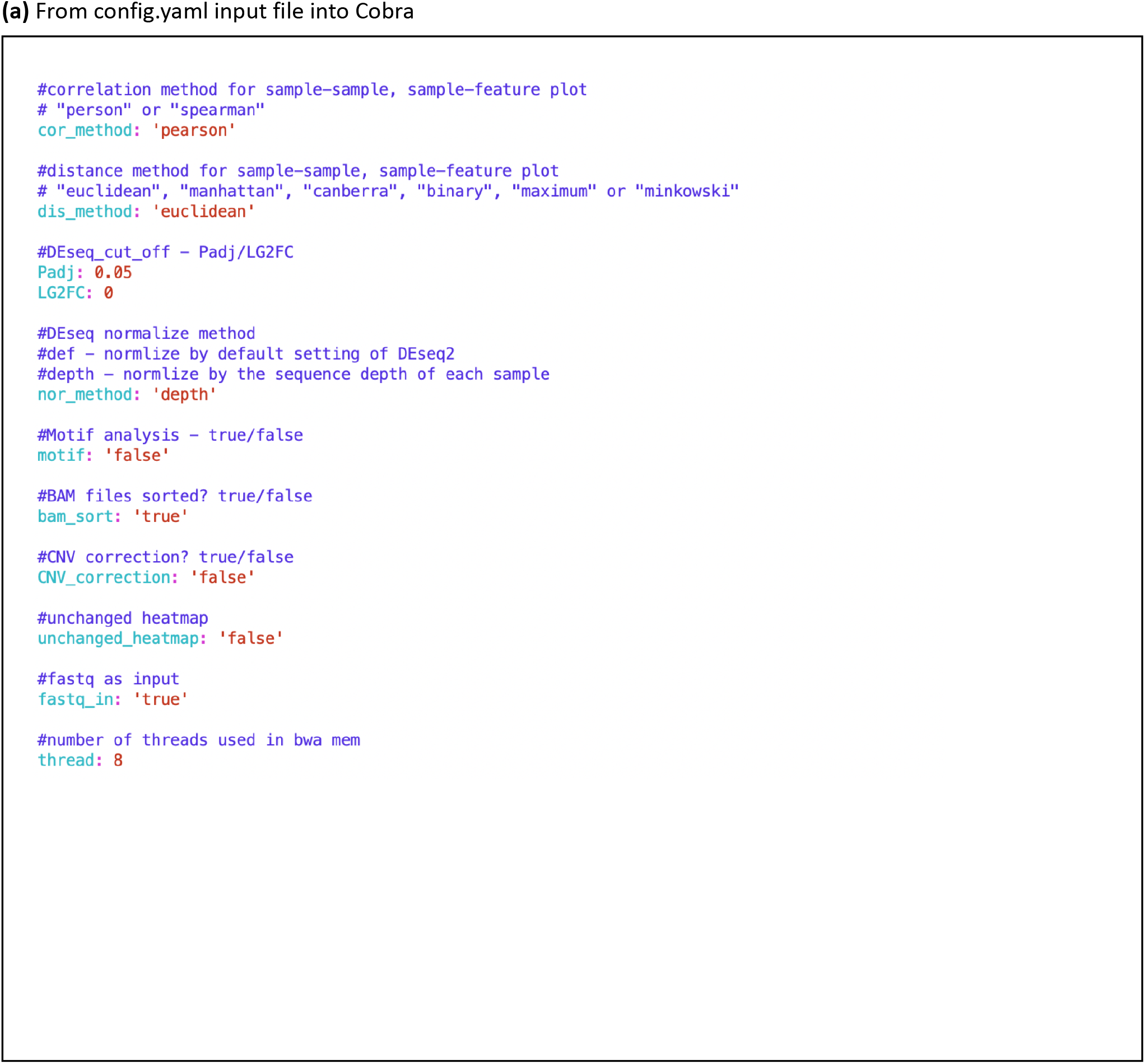
b) Example of parameter setup in config.yaml file.

**Supplementary Figure 2c.**
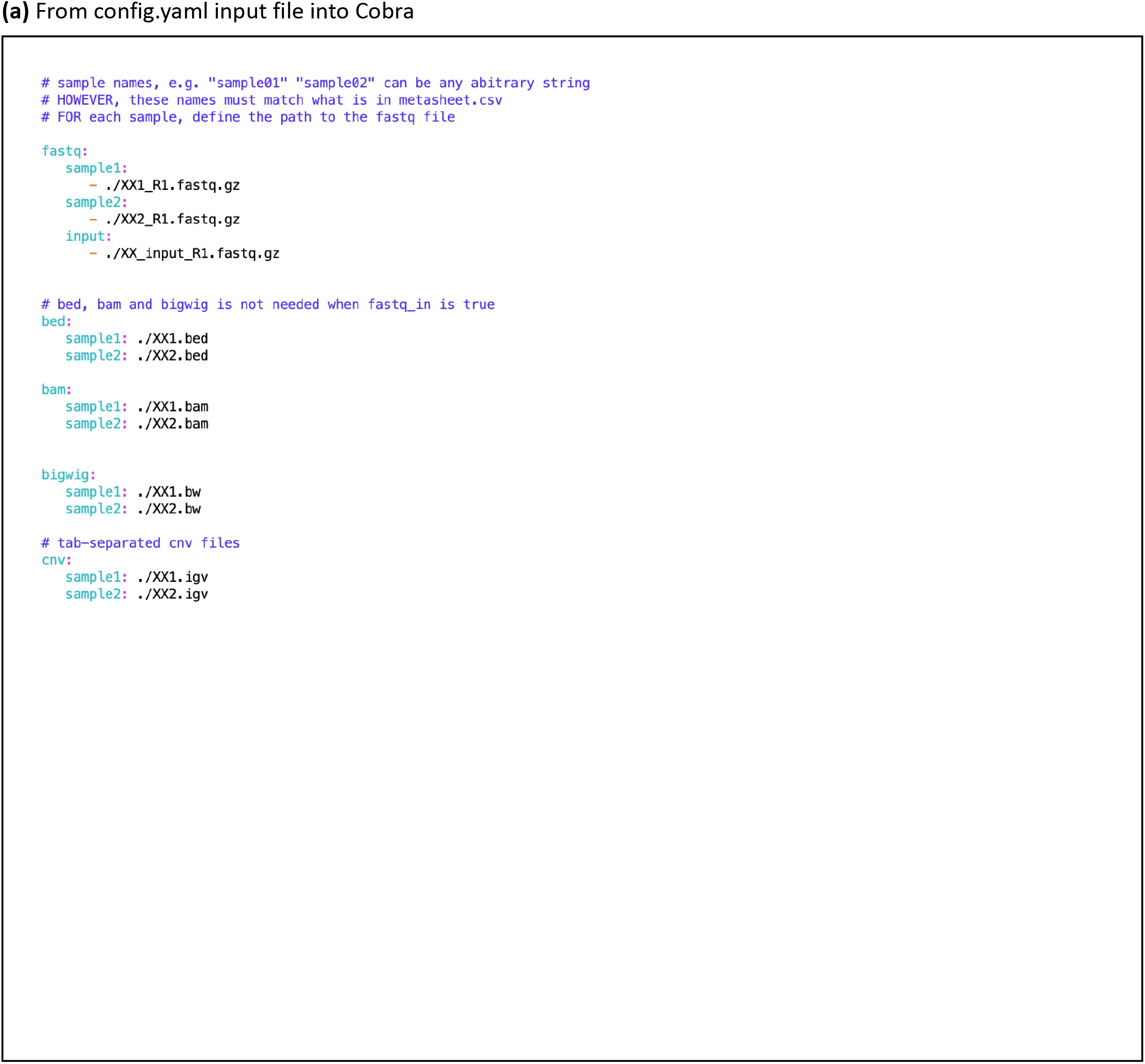
c) Example of sample path setup in config.yaml file.

**Supplementary Figure 3.**
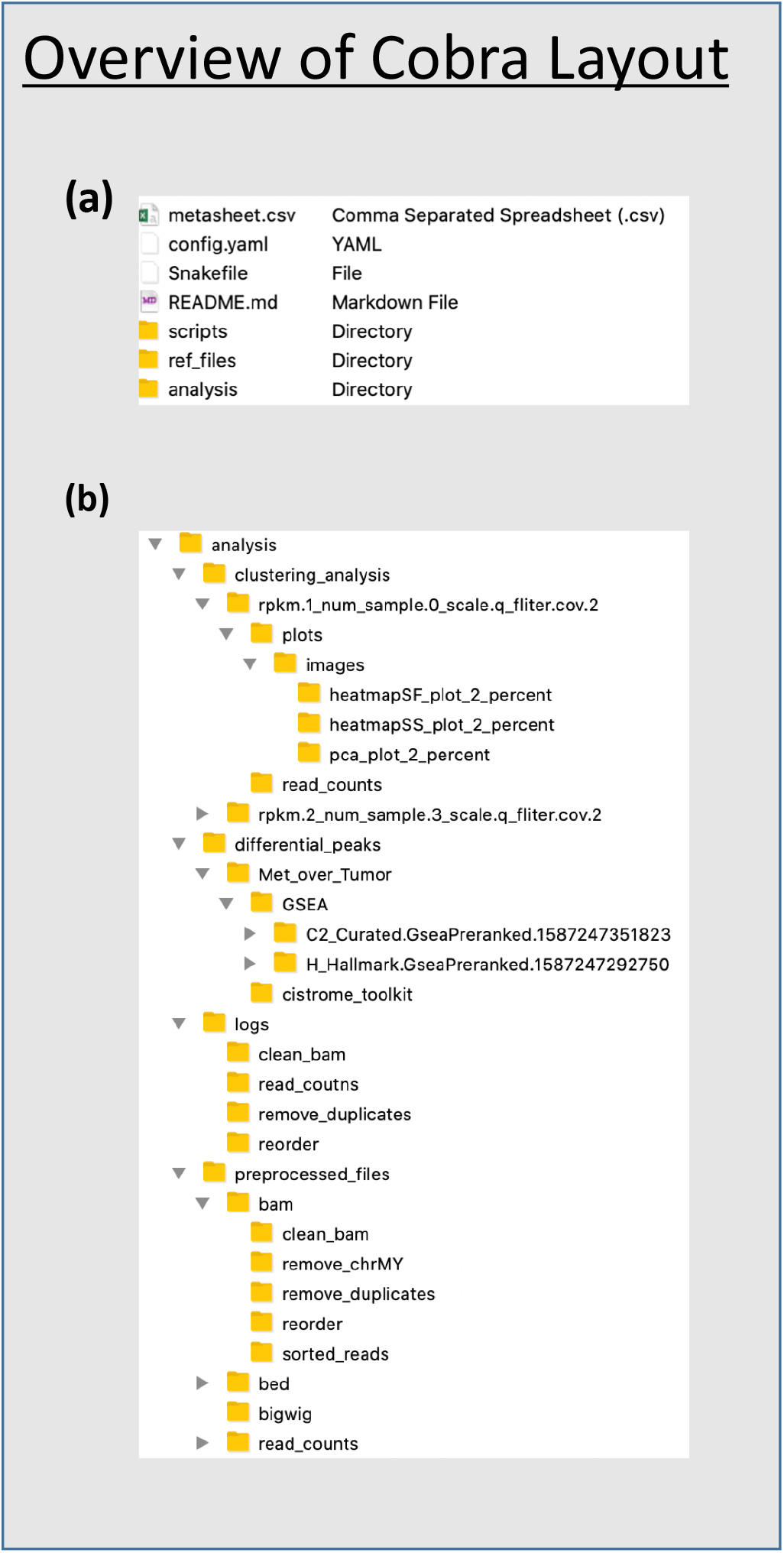
File structure of CoBRA input and output.

## Notes

### Competing Interest Statement

The authors have declared no competing interest.

